# Shared and Specific miRNA-mRNA in Human and Murine Acute Kidney Injury

**DOI:** 10.64898/2026.10.30.762662

**Authors:** Danielle Janosevic, Thomas W. McCarthy, Ashish Sadolikar, Takashi Hato, Michael T. Eadon, Pierre Dagher, the Kidney Precision Medicine Project

**Affiliations:** Division of Nephrology, Indiana University School of Medicine, Indianapolis, IN 46202 USA

**Author notes:** Corresponding information: Danielle Janosevic, Indiana University School of Medicine, Department of Medicine, 950 W. Walnut St, Research 2, Indianapolis, IN 46202.

**Keywords:** acute kidney injury, cecal ligation and puncture, ischemia-reperfusion injury, lipopolysaccharide, microRNA, cross-species transcriptomics, human AKI

## Abstract

Acute kidney injury (AKI) is managed as a single clinical entity despite substantial biological heterogeneity. We established a cross-species framework to define etiology-specific molecular programs and determine how these programs relate to molecular heterogeneity and pathobiology in human AKI. We performed parallel sequencing of totalRNA (mRNA) and miRNA (kidney, whole blood) in 3 murine models (mild AKI: ischemia-reperfusion injury, IRI; cecal-ligation and puncture, CLP; endotoxemia, LPS); kidney mRNA and miRNA clearly distinguished the models, blood mRNA maintained model-specific separation, and blood miRNA was less distinct. IRI was characterized by epithelial injury/repair, and extracellular matrix remodeling (i.e., *miR-125b-1-3p, Vim, Sprr2g*); CLP, broader inflammatory, metabolic programs (i.e., *miR-692, Pfkfb3, Cdo1*); LPS, inflammatory signaling (i.e., *miR-155-5p, Irf7, Gbp3*). We then compared these signatures to human whole blood RNAseq. Human blood samples varied along a shared gene axis that correlated to differing degrees with all three murine models, consistent with the heterogeneous nature of clinical AKI rather than discrete assignment to a single experimental etiology. Orthologs were linked to immune processes (i.e., *SIGLEC14, DAAM2, CD177, DEFA3*), and the presence of tubular casts in human biopsies, reflecting that mild murine AKI models capture etiology-specific biology relevant to severe human AKI. Integration of injury etiology, severity, and time may provide a framework for developing circulating biomarkers that may resolve the biological heterogeneity of human AKI.

## Introduction

Acute kidney injury (AKI) is clinically defined by changes in serum creatinine and urine output, yet these criteria encompass biologically diverse processes within a single syndrome.^1; 2^ Distinct etiologies, including ischemic injury (ischemia-reperfusion injury models, IRI), abdominal polymicrobial sepsis (cecal ligation and puncture, CLP), and endotoxemia (lipopolysaccharide administration, LPS), differ in their initiating mechanisms, systemic context, and temporal evolution, but ultimately converge on common pathophysiologic features such as tubular epithelial injury, microvascular dysfunction, immune activation, and metabolic stress.^2–6^

Human single-cell and spatial transcriptomic studies, and earlier foundational studies, further demonstrate that clinically defined AKI comprises diverse epithelial, immune, and stromal cell states rather than a uniform molecular phenotype.^7–21^ Experimental models offer an opportunity to dissect this heterogeneity by etiology.^3; 5; 12; 13; 22^ IRI directly disrupts renal perfusion and reperfusion, causing renal injury.^23; 24^ CLP models polymicrobial abdominal sepsis, with injury severity influenced by procedural parameters and supportive care.^25; 26^ In contrast, LPS induces endotoxemia without a replicating microbial source, eliciting a rapid, reversible inflammatory response with dose- and duration-dependent renal effects.^5; 6; 27–29^ For all three models, we chose mild forms of acute kidney injury recapitulated in each model, which we hypothesized would help resolve distinct early initiating factors rather than common AKI sequelae.

Integrated miRNA and mRNA analysis provides a complementary regulatory perspective because miRNAs can coordinately modulate networks of target genes and often exhibit distinct expression patterns in kidney tissue and blood compartments.^30–34^ However, most studies have focused on individual AKI models or a single RNA modality, limiting assessment of whether miRNA-mRNA regulatory relationships are conserved or etiology-specific.^17; 35–37^ We hypothesized that, although these models share components of a common renal injury response, their distinct initiating mechanisms generate etiology-specific molecular programs in the kidney and blood compartments that could inform future precision therapeutic strategies. To test this hypothesis, we performed paired miRNA and total RNA sequencing of kidney and whole blood across murine IRI, CLP, and LPS using milder injury conditions designed to capture early molecular phenotypes of AKI. We further hypothesized that etiology-associated murine signatures are detectable in blood and conserved in human AKI. By mapping murine blood signatures to human transcriptomes, we sought to determine whether these molecular programs resolve biological heterogeneity among patients with AKI and provide a framework for mechanism-based patient stratification.

## Results

### Murine AKI models demonstrate distinct kidney mRNA profiles despite shared tubular injury markers

Three mild injury models were leveraged to uncover distinct molecular profiles: IRI (9 minutes), CLP (1 puncture), and LPS (2.5 mg/kg i.v.). Kidney injury transcriptomic markers *Havcr1* and *Lcn2* increased in CLP, IRI, and LPS relative to matched controls, confirming an intrarenal molecular injury pattern in all three models (Figure 1A). Serum creatinine values were increased in IRI and CLP, but not the LPS model (Figure 1B). Pathologic tubular scoring was performed in all three models relative to controls, with a similar pattern of acute tubular injury (ATI) noted (Figure 1E). The combination of injury transcript induction without functional decline suggests that the LPS model was relatively less severe at this dosage compared to CLP or IRI.

**Figure 1.**
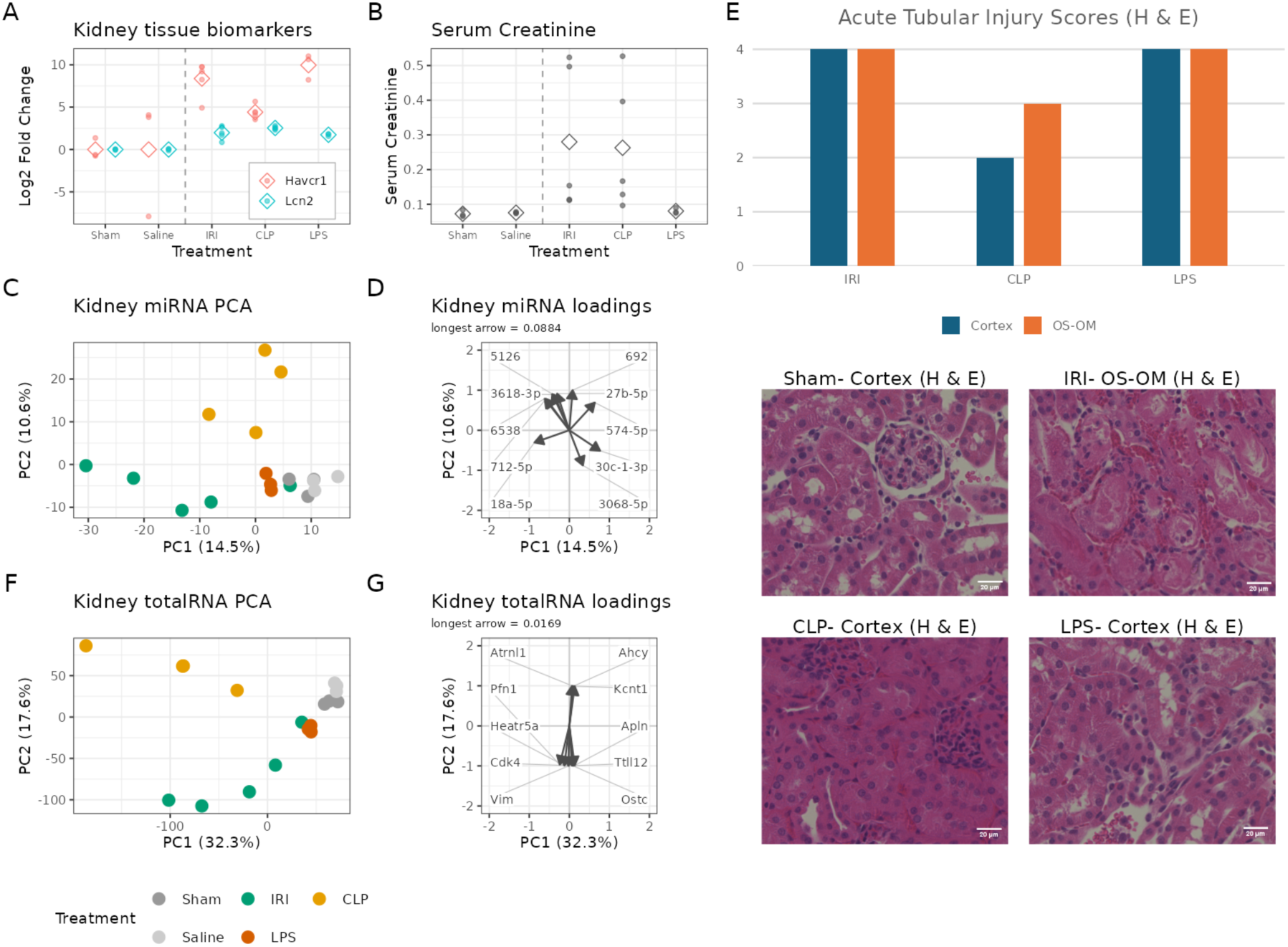
Phenotypic and molecular characterization of experimental AKI models. (A) RT-qPCR analysis of kidney tubular injury markers *Havcr1* and *Lcn2* across experimental AKI models relative to Sham and Saline controls. Points represent individual samples; diamonds indicate group means. (B) Serum creatinine (mg/dL) concentrations across experimental groups. Points represent individual animals; diamonds indicate group means. (C) Principal component analysis (PCA) of kidney miRNA expression. Each point represents an individual sample and is colored by treatment. Variance explained by each principal component (PC) is indicated in parentheses. (D) Loadings of the top 10 kidney miRNAs by loading weight on PC1 and PC2. Arrows are scaled relative to the longest loading vector, with the raw loading value of the longest vector indicated above the panel; the raw loading of each remaining vector corresponds to its relative arrow length multiplied by this value. (E) Acute tubular injury scores for IRI, CLP, and LPS (cortex and OS-OM) and representative H&E-stained kidney sections (Sham, IRI, CLP, and LPS). Bars represent the averaged injury score (see methods) from a representative mouse for each model. Sections were imaged using a 20× objective and enlarged for visualization. (F) PCA of kidney total RNA expression. Each point represents an individual sample and is colored by treatment. Variance explained by each PC is indicated in parentheses. (G) Loadings of the top 10 kidney total RNA genes by loading weight on PC1 and PC2, scaled as described in (D). Treatment colors shown below (F) apply to (C) and (F). Sample sizes were n = 5 for IRI and n = 4 for CLP, except serum creatinine, which was available for n = 5 CLP animals; n = 3 per Sham, Saline, and LPS group. H&E, hematoxylin and eosin; IRI, ischemia-reperfusion injury; CLP, cecal ligation and puncture; LPS, lipopolysaccharide; PCA, principal component analysis; PC, principal component; OS-OM, outer stripe of the outer medulla.

Although tubular injury markers were induced across IRI, CLP, and LPS, including other injury transcripts such as *Timp1, Hmox1* (Supplemental Figure 1), their miRNA and mRNA responses differed substantially. Principal component analysis (PCA) of mRNA and miRNA expression demonstrated distinct clustering by injury model (Figure 1C,D,F,G). Both miRNA and mRNA profiles separated injured samples from controls along PC1, while PC2 distinguished CLP from IRI and LPS. We used miRNA and mRNA loadings to determine the key transcripts that drove PCA clustering of the AKI models (Figure 1D, G). Dominant kidney miRNA loadings, separating controls from IRI and CLP along PC1, included miR-18a-5p and miR-3068-5p, while miR-692 and miR-712-5p contributed to separation between CLP and IRI models. Kidney mRNA separation was driven in part by *Pfn1* and *Vim* along PC1 and by *Ahcy* and *Kcnt1* along PC2. Collectively, these loadings identified cytoskeletal remodeling (*Pfn1* and *Vim*) along PC1, and epithelial state change/function (*Kcnt1*), stress adaptation (*Ahcy*) along PC2.

We then compared the similarities and differences of the injury response across IRI, CLP, and LPS models. Differential-expression and UpSet plot analysis demonstrated that CLP generated the largest kidney response: 1,957 genes were upregulated and 1,959 were downregulated. IRI had 897 upregulated and 663 downregulated genes, whereas LPS had 453 upregulated and 165 downregulated genes. Only 91 upregulated genes and 48 downregulated genes were shared across IRI, CLP, and LPS, while the largest intersections were model-specific: 1,460 upregulated and 1,533 downregulated genes were unique to CLP; 425 and 243 were unique to IRI; and 243 and 73 were unique to LPS (Figure 2B-C; Supplemental Figure 2A, C-D, Supplemental Tables 1,2).

**Figure 2.**
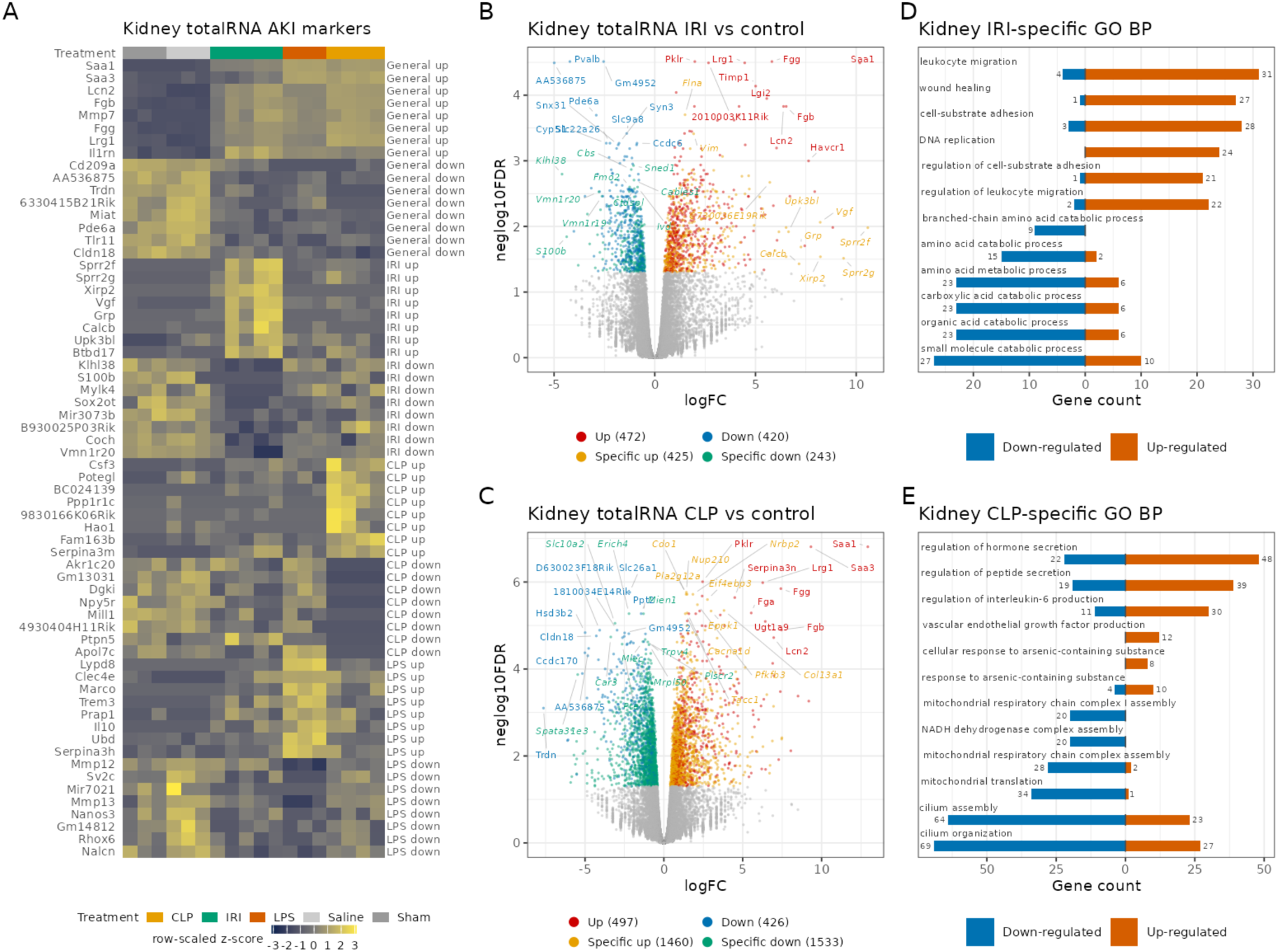
Shared and model-specific kidney mRNA programs across murine AKI models. (A) Heatmap of kidney total RNA genes shared across AKI models and treatment-specific marker genes, ordered by treatment group. Expression values are shown as row-scaled z scores. (B) Volcano plot of kidney total RNA differential expression in IRI versus control. (C) Volcano plot of kidney total RNA differential expression in CLP versus control. In (B) and (C), genes are classified as upregulated, downregulated, treatment-specific upregulated, or treatment-specific downregulated as indicated. (D) Gene Ontology (GO) Biological Process terms enriched among IRI-specific kidney total RNA differentially expressed genes (DEGs), shown separately for upregulated and downregulated genes. (E) GO Biological Process terms enriched among CLP-specific kidney total RNA DEGs, shown separately for upregulated and downregulated genes. Differentially expressed genes were defined by FDR < 0.05 and absolute fold change ≥ 1.2. Heatmaps were generated from log2 counts per million (CPM) values, with treatment-specific and shared features ranked by absolute fold change and standardized to z scores where indicated. GO Biological Process enrichment was performed (Benjamini-Hochberg correction, P < 0.05, q < 0.20), using all genes tested in the corresponding contrast as the background. n = 5 for IRI, n = 4 for CLP, and n = 3 for Sham, Saline, and LPS. IRI, ischemia-reperfusion injury; CLP, cecal ligation and puncture; LPS, lipopolysaccharide; GO, Gene Ontology; DEG, differentially expressed gene; FDR, false discovery rate; CPM, counts per million.

Our Gene Ontology biological process enrichment analysis further highlighted the divergence between injury phenotypes. IRI-specific upregulated genes *Sprr2f/g*, related to epithelial cell barrier function and also enriched in pathways related to response to regulation of cell adhesion, leukocyte migration, and wound healing, whereas IRI-specific downregulated genes (e.g., *Cbs*) were enriched for lipid, amino-acid, monocarboxylic-acid, and related metabolic or catabolic processes (Figure 2B, D, Supplemental Table 3). In contrast, CLP-specific upregulated genes included *Pfkfb3*, with enrichment observed for regulation of secretion, responses to peptide or hormone signals, peptide transport, vesicle-mediated transport and regulation of interleukin-6 production. CLP-specific downregulated pathways included cilium organization and assembly, mitochondrial processes and related biosynthetic programs, reflecting the broader systemic effects of CLP (Figure 2C,E, Supplemental Table 3). LPS generated a kidney gene-expression response with a narrow mRNA signature. LPS-specific upregulated genes (e.g., *Lypd8, Irf7*) were enriched for response to virus, regulation and activation of innate immune responses, defense responses, and interferon-beta signaling (Supplemental Figure 2A-B, Supplemental Table 3). Thus, LPS supports select innate and interferon-associated components relevant to CLP injury, but fails to fully recapitulate the CLP injury program.

### Murine kidney miRNAs reflect etiology- and cell type-specific regulatory programs

We next assessed the differentially expressed kidney miRNAs. We identified eight upregulated miRNAs in IRI, nine in CLP, and five in LPS (Figure 3A-D, Supplemental Table 4). UpSet plots revealed limited overlap, with only a single upregulated miRNA, miR-132-5p, shared across all three models (Figure 3A, Supplemental Figure 3A, Supplemental Table 2). Two unique downregulated miRNAs were detected in CLP, while no downregulated kidney miRNAs met the applied threshold in IRI or LPS (Supplemental Figure 3A-B). miR-223-3p and miR-125b-1-3p were unique to IRI; miR-692, miR-92a-1-5p, miR-1247-3p, miR-27a-5p, and miR-27b-5p were unique to CLP; miR-503-5p and miR-671-5p were downregulated in CLP; and miR-155-5p and miR-132-3p were unique to LPS (Figure 3A).

**Figure 3.**
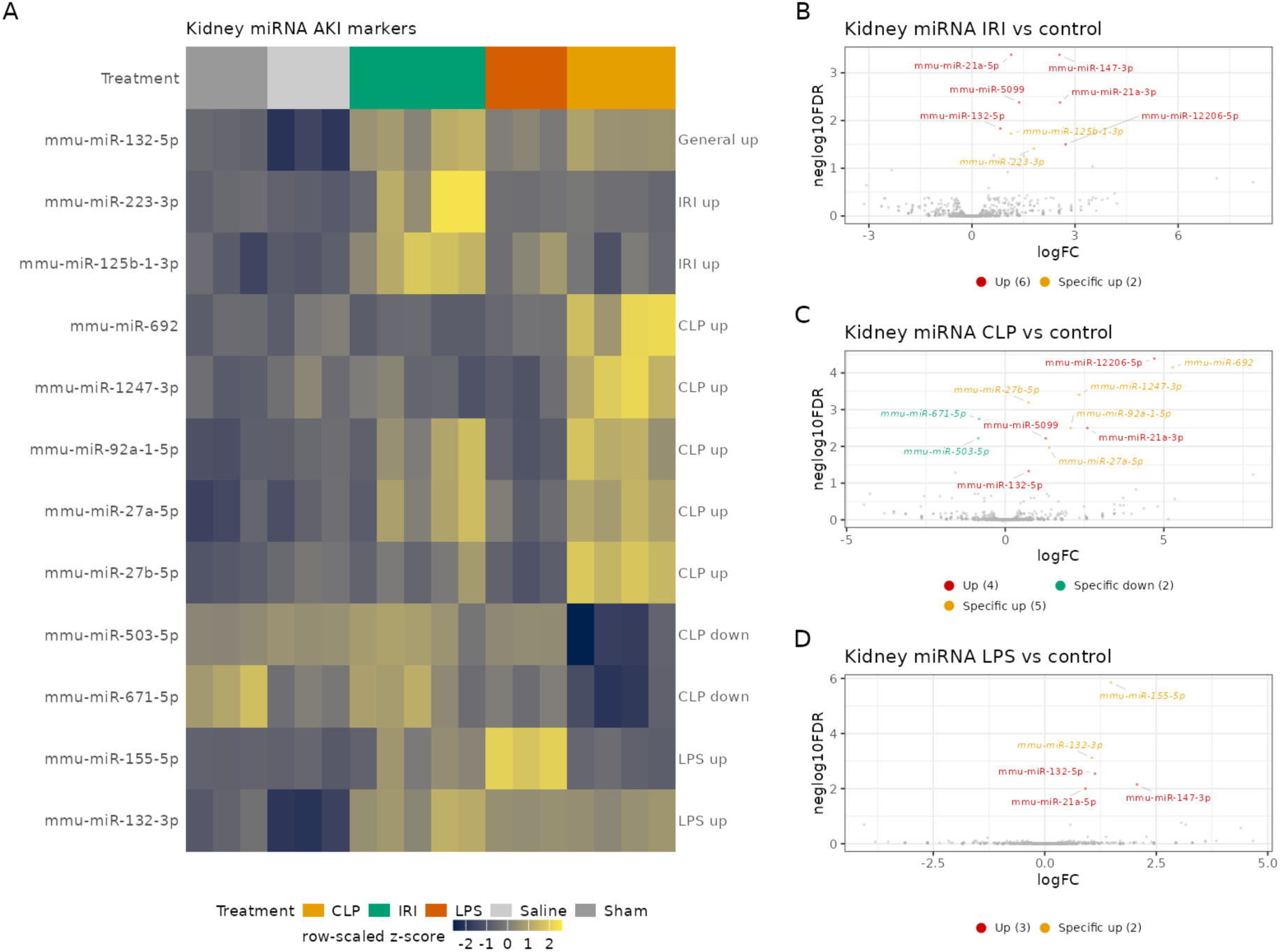
Shared and model-specific kidney miRNA expression responses across murine AKI models. (A) Heatmap of kidney miRNAs shared across AKI models and treatment-specific miRNAs, ordered by treatment group. Expression is shown as row-scaled z scores. (B–D) Volcano plots of kidney miRNA differential expression in IRI (B), CLP (C), and LPS (D) versus control. Differentially expressed miRNAs are classified as upregulated, downregulated, or treatment-specific as indicated. Differentially expressed miRNAs were defined as FDR < 0.05 and |logFC| ≥ log2(1.2). IRI, ischemia-reperfusion injury; CLP, cecal ligation and puncture; LPS, lipopolysaccharide; FDR, false discovery rate.

We next constructed a single-cell atlas derived from these models (Figure 4A-B). Representative model-specific miRNA target scores were projected onto the single-cell UMAP to infer cell-type specificity in injury. We projected the top unique miRNA for each model, including: IRI-associated miR-223-3p, CLP-associated miR-692, and LPS-associated miR-155-5p (Figure 4C-H). The IRI-related miR-223-3p target program was identified in epithelial clusters of the proximal tubule (PT) and collecting duct (CD), with mapping to the endothelial and stromal compartments, but had too few target genes for enrichment (Figure 4F). For CLP, the miR-692 target program was identified broadly across epithelial, immune, and endothelial clusters, with downstream genes enriched in metabolic-related pathways (Supplemental Figure 4B, Supplemental Table 5). In contrast, the LPS-specific miR-155-5p target program was concentrated in the endothelial and immune clusters (Figure 4C-E), with target gene enrichment in immune and defense response-related pathways (Supplemental Figure 4C, Supplemental Table 5). These results indicate that AKI etiologies engage distinct downstream gene programs within various cell types.

**Figure 4.**
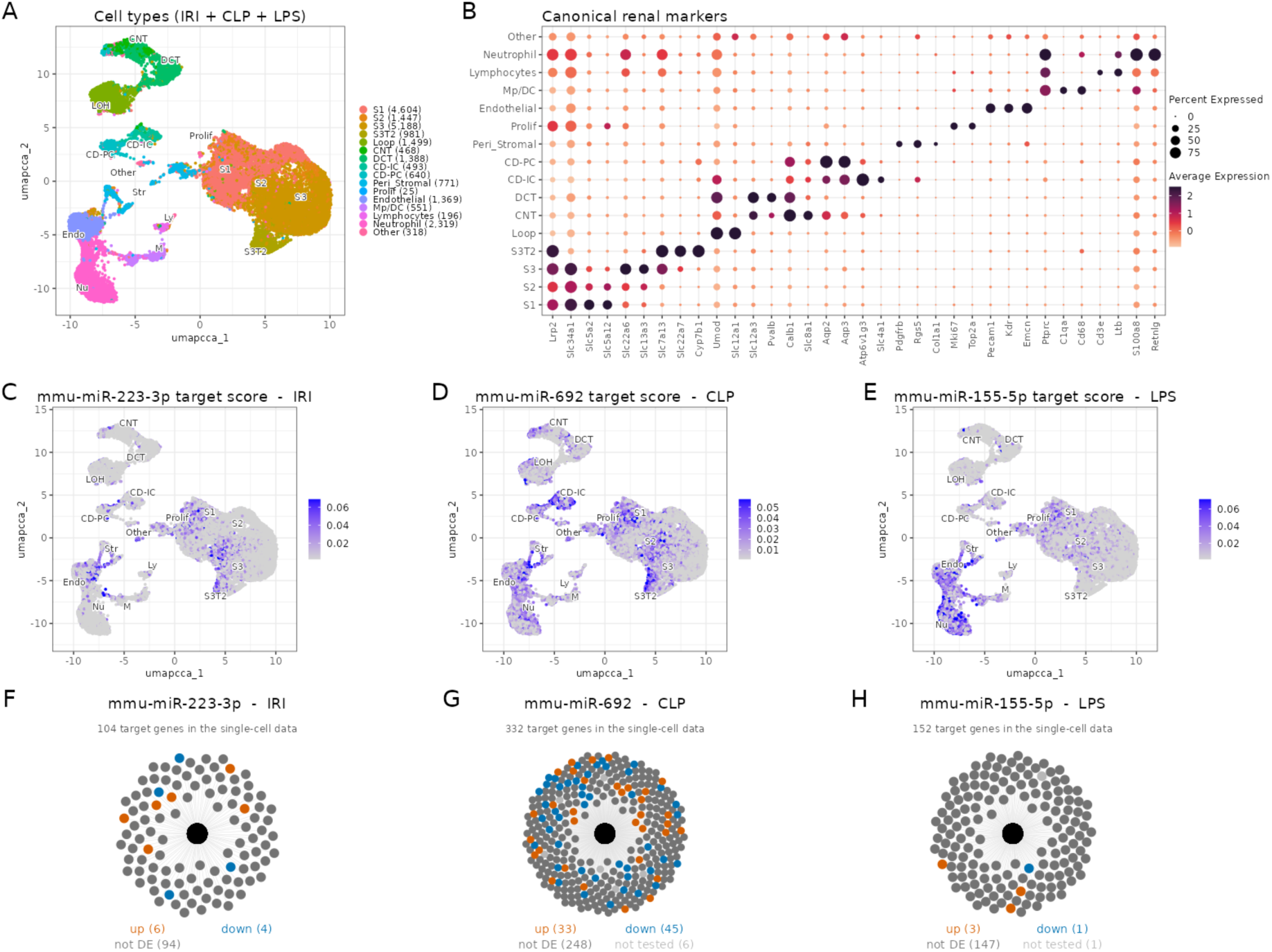
Cell type distribution and predicted miRNA target expression across experimental AKI models. (A) UMAP of integrated single-cell kidney data from IRI, CLP, and LPS models, colored by cell type. Cell-type labels shown in (A) also apply to (C–E). (B) Expression of canonical renal marker genes across cell types. Dot size indicates the percentage of cells expressing each gene, and color indicates average expression. (C–E) UMAPs showing miRDB target module scores for mmu-miR-223-3p in IRI (C), mmu-miR-692 in CLP (D), and mmu-miR-155-5p in LPS (E). (F–H) Predicted target genes for mmu-miR-223-3p (F), mmu-miR-692 (G), and mmu-miR-155-5p (H), colored according to differential expression in the corresponding kidney total RNA comparison. Orange indicates upregulated, blue indicates downregulated, dark gray indicates not differentially expressed, and light gray indicates not tested. The central black node represents the miRNA; node positions carry no quantitative meaning. IRI, ischemia-reperfusion injury; CLP, cecal ligation and puncture; LPS, lipopolysaccharide; UMAP, uniform manifold approximation and projection.

### Murine blood mRNA preserves etiology-specific molecular programs

Whole-blood mRNA clearly distinguished injury models (Figure 5A). LPS and CLP, to a lesser degree, showed the largest separation from saline controls, whereas IRI clustered more closely to sham controls. Genes with positive PC1 mRNA loadings, including *Wdtc1*, *Myh9*, *Actn1,* were enriched for chromosome segregation, cell-cycle regulation, cytokinesis, and mitotic spindle assembly. Negative PC2 loadings, including *Gbp3*, showed more limited enrichment for T-helper 1 cell cytokine production and proteasome assembly (Figure 5B; Supplemental Table 6). Thus, whole blood retained model-specific molecular information, although the dominant circulating programs differed from those observed in kidney tissue.

**Figure 5.**
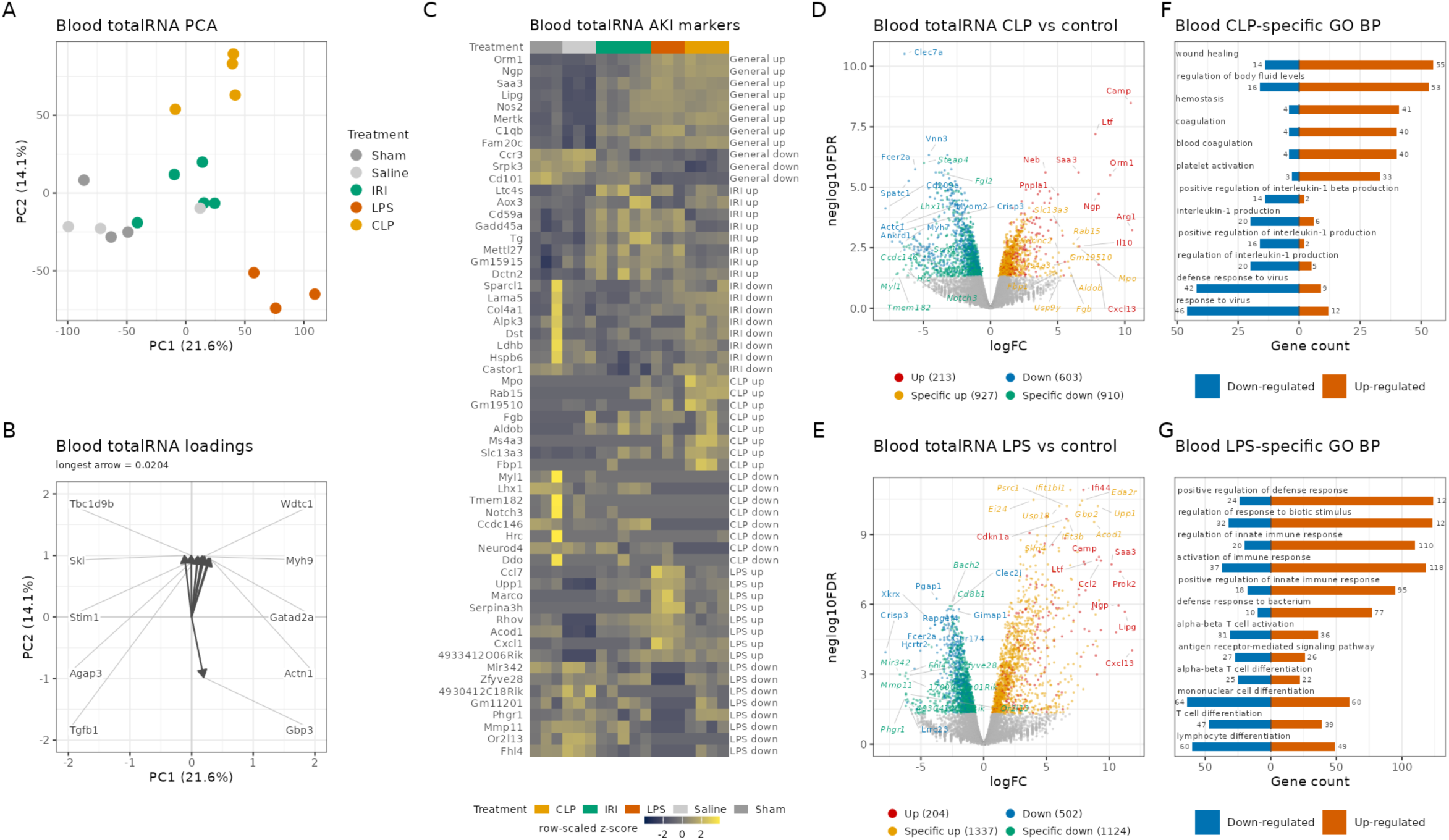
Shared and model-specific blood gene expression programs across murine AKI models. (A) PCA of blood total gene expression (PC1 vs PC2). Each point represents one sample, colored by treatment. Variance explained by each principal component (PC) is indicated in parentheses. (B) Top 10 genes by loading weight on PC1 and PC2. Arrows are scaled relative to the longest loading vector, with the raw loading value of the longest vector indicated above the panel; the raw loading of each remaining vector corresponds to its relative arrow length multiplied by this value. (C) Heatmap of genes shared across AKI models and treatment-specific marker genes, ordered by treatment group. Expression is shown as row-scaled z scores. (D and E) Volcano plots of blood total RNA differential expression in CLP (D) and LPS (E) versus control. Genes are classified as upregulated, downregulated, or treatment-specific as indicated. (F and G) GO Biological Process terms enriched among CLP-specific (F) and LPS-specific (G) DEGs, shown separately for upregulated and downregulated genes. DEGs were defined as previously described in Figure 2. IRI, ischemia-reperfusion injury; CLP, cecal ligation and puncture; LPS, lipopolysaccharide; GO, Gene Ontology; DEG, differentially expressed gene; PC, Principal Component; PCA, Principal Component Analysis.

Differential expression and UpSet analyses further demonstrated etiology-specific separation in the blood compartment (Figure 5D-E, Supplemental Figure 5A, C-D, Supplemental Tables 1,2). LPS had 1,541 upregulated and 1,626 downregulated blood genes, CLP had 1,140 upregulated and 1,513 downregulated genes, and IRI had 41 upregulated and 118 downregulated genes. Of these genes, 11 upregulated genes were shared between the three models (e.g., *Saa3, Nos2, C1qb, Spp1*), primarily related to a systemic inflammatory response (Figure 5C, D,E; Supplemental Figure 5A).^38^ CLP-specific upregulated genes were enriched for platelet activation hemostasis and coagulation (*Fgb*) and neutrophil function (*Mpo*). CLP-specific downregulated genes were enriched for antiviral responses, regulation of interleukin-1 production, innate immune activation, and genes related to endoplasmic reticulum stress (i.e., *Hrc*, Figure 5D,F, Supplemental Table 3). LPS-specific upregulated pathways showed broad enrichment for innate immune activation and pattern-recognition receptor signaling (*Marco*), defense responses to bacteria and viruses (*Ccl7*), cytokine-mediated signaling, inflammatory regulation, and response to lipopolysaccharide (*Cxcl1*, Figure 5E,G, Supplemental Table 3).^38^ LPS-specific downregulated pathways were enriched for lymphocyte, mononuclear-cell activation (Figure 5G, Supplemental Table 3). The modest IRI-specific upregulated gene set was enriched for terms related to negative regulation of blood vessel and angiogenesis pathways (e.g., *Gadd45a*, see Supplemental Figure 5A-B).^38^ Together, the blood mRNA data distinguished a platelet and hemostatic program in CLP, a broad innate inflammatory program in LPS, and a comparatively limited circulating response in IRI characterized by the negative regulation of angiogenesis.

### Murine blood miRNAs exhibit a broader response to AKI

The blood miRNA profiles were able to distinguish control from AKI samples, although the degree of separation between models was more modest (Figure 6A). LPS produced by far the largest set of model-specific miRNA changes with 105 upregulated and 112 downregulated miRNAs, whereas CLP and IRI showed more restricted signatures, with less than 20 differentially expressed miRNAs in total (Supplemental Figure 6A-B, Supplemental Table 4), and no miRNA shared between all 3 models.

**Figure 6.**
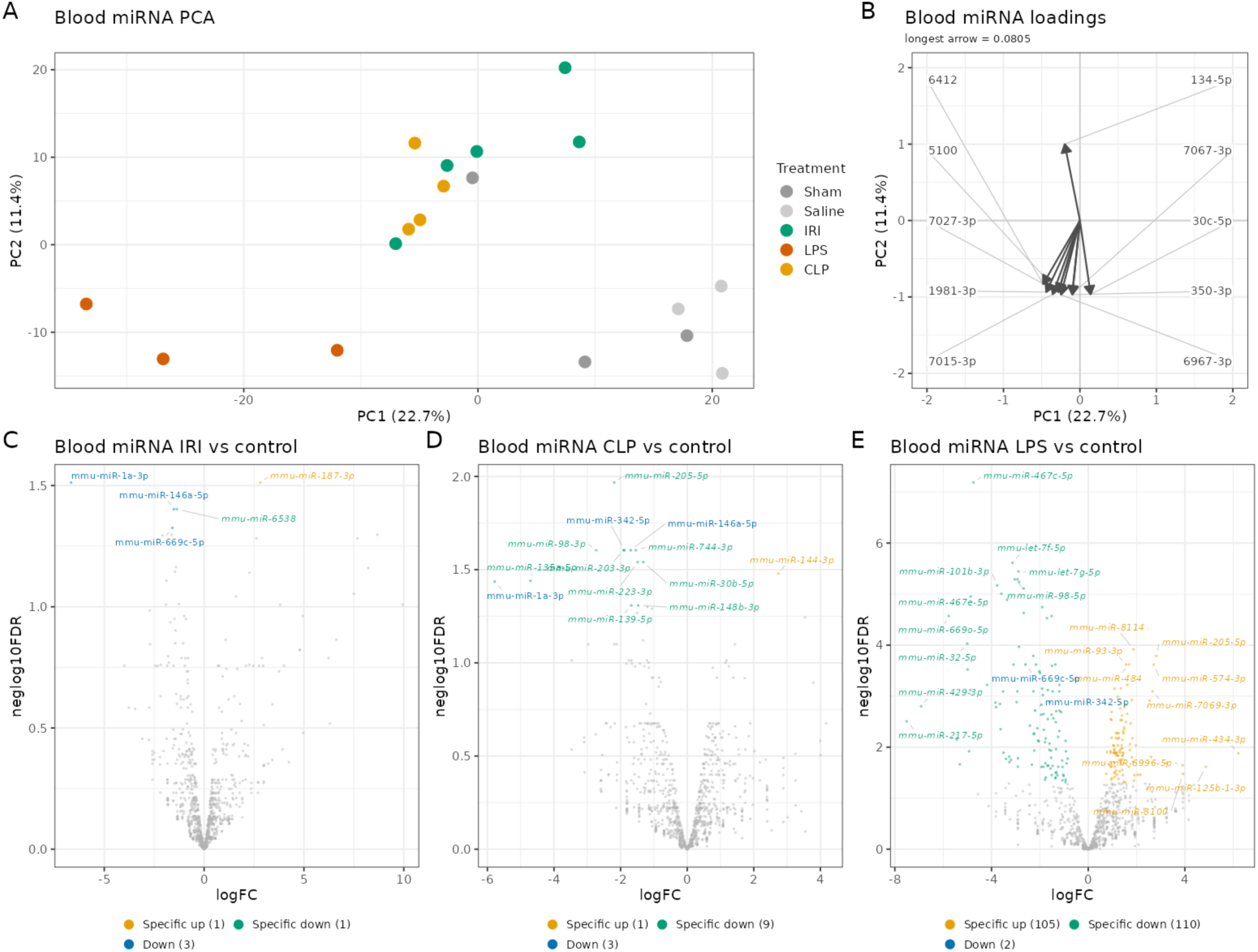
Blood miRNA profiles and differential expression across murine AKI models. (A) Principal component analysis (PCA) of blood miRNA expression. Each point represents an individual sample and is colored by treatment; variance explained by each principal component (PC) is indicated in parentheses. (B) Loadings of the top 10 blood miRNAs by loading weight on PC1 and PC2. Arrows are scaled relative to the longest loading vector, with its raw loading value indicated above the panel. (C–E) Volcano plots of blood miRNA differential expression in IRI (C), CLP (D), and LPS (E) versus control. Differentially expressed miRNAs are classified as upregulated, downregulated, or treatment-specific as indicated. Differentially expressed miRNAs were defined as previously described in Figure 3. IRI, ischemia-reperfusion injury; CLP, cecal ligation and puncture; LPS, lipopolysaccharide; PCA, principal component analysis; PC, principal component; FDR, false discovery rate.

We compared the blood miRNA signature between AKI models. CLP was characterized by increased miR-144-3p and model-specific decreases in miR-205-5p, miR-98-3p, miR-223-3p, and miR-30 family members, many of which are involved in epithelial cell survival and proliferation.^39–41^ IRI-specific increases included miR-187-3p, serving a protective role in the renal glomerular filtration barrier.^42^ LPS showed broad miRNA-related regulation, including increased miR-93-3p, miR-205-5p, and miR-574-3p and decreased miR-467e-5p, miR-98-5p, many of which are involved in regulating the inflammatory response, epithelial-to-mesenchymal transition, and apoptosis.^41; 43^ Notably, miR-205-5p changed in opposite directions in CLP and LPS, further supporting distinct circulating regulatory responses between these two infection-related models (Figure 6D-E). The strong circulating LPS signature despite mild renal functional injury highlights the contribution of systemic inflammation to blood RNA profiles. These data indicate a relatively limited systemic inflammatory response, particularly in mild IRI, when compared to renal tissue miRNA profiles in our study.^44^

### Human and Murine blood signatures of AKI have overlapping molecular profiles

We then investigated whether etiology-specific murine signatures are detectable in the blood of human subjects with AKI. We compared Spearman correlations of human blood samples with murine IRI, CLP, and LPS blood total RNA. We identified a dominant shared pattern in all murine models; samples with stronger positive correlation to one murine injury program generally showed concordant positive correlation with the other murine AKI models, whereas samples at the opposite end of the continuum were negatively correlated across murine AKI models. Interestingly, human samples with dominant cast features on histopathology, a feature of significant renal damage, were most correlated with the murine mild-injury models (Figure 7C, Supplemental Table 7). Notably, the cross-species correlation patterns were not clustered by KDIGO stage (Figure 7C). These findings are consistent with a human blood transcriptome shaped by overlapping injury mechanisms, systemic host responses, comorbid conditions, treatment exposures, and circulating cell composition rather than by a single isolated experimental insult.

**Figure 7.**
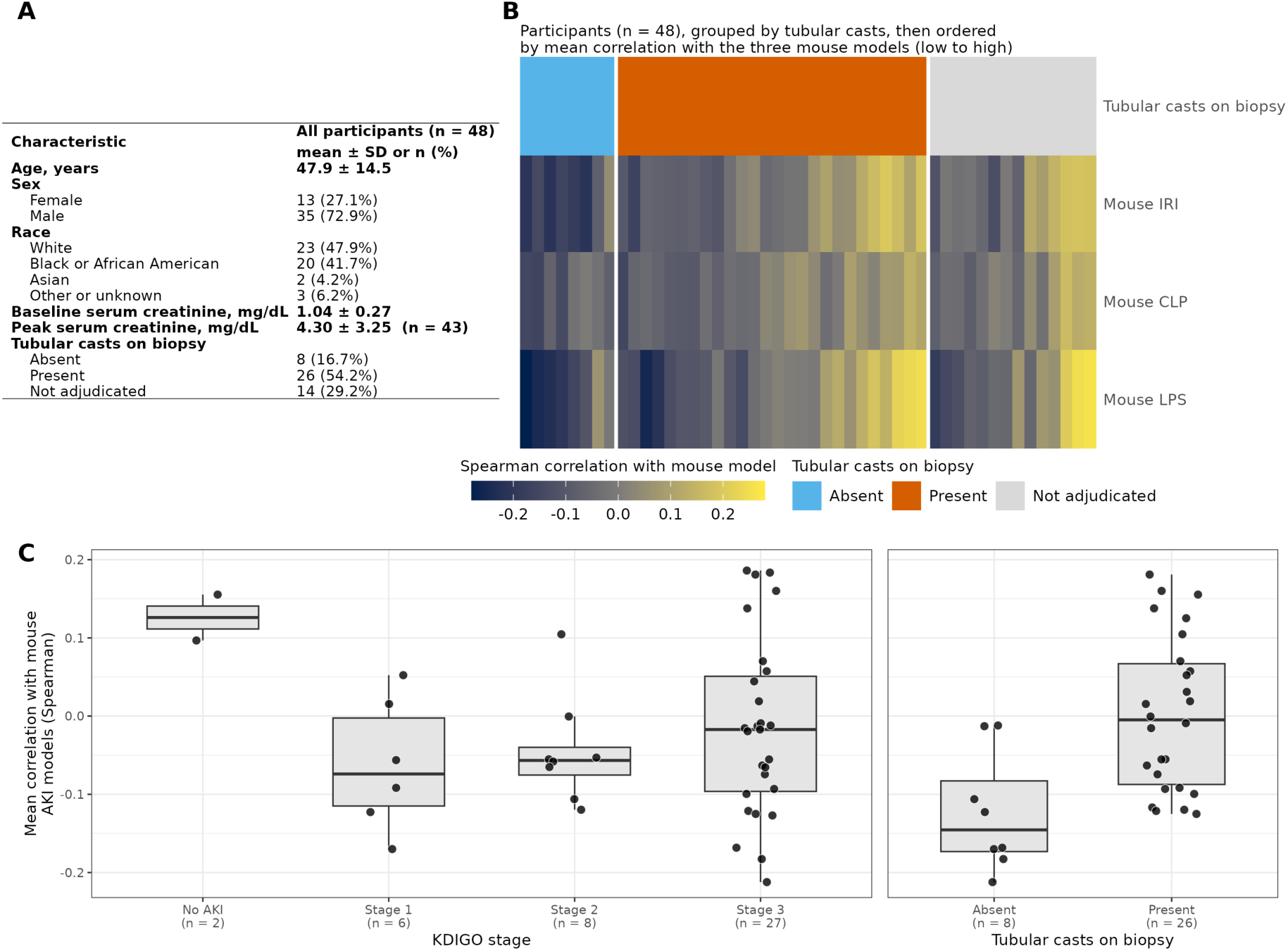
Cross-species similarity of human AKI blood transcriptomes to experimental mouse AKI models. (A) Demographic and clinical characteristics of 48 KPMP participants in the AKI cohort. Continuous variables are mean ± SD and categorical variables are n (%). (B) Spearman correlations between individual participant whole-blood transcriptomes and mouse blood transcriptional responses to IRI, CLP, and LPS across 10,568 one-to-one orthologous genes. Gene expression was sex-adjusted and centered across the cohort. Participants are grouped by tubular casts on biopsy and ordered within groups by mean correlation across the three mouse models. (C) Mean correlation across the three mouse models by KDIGO stage (No AKI, n = 2; Stage 1, n = 6; Stage 2, n = 8; Stage 3, n = 27) and tubular casts on biopsy (absent, n = 8; present, n = 26). Boxes indicate median and interquartile range, and points represent individual participants. Five participants without KDIGO staging and 14 without biopsy adjudication are excluded from the corresponding panels. No statistical comparisons were performed. KPMP, Kidney Precision Medicine Project; AKI, acute kidney injury; IRI, ischemia-reperfusion injury; CLP, cecal ligation and puncture; LPS, lipopolysaccharide; KDIGO, Kidney Disease: Improving Global Outcomes.

Human total-RNA principal-component analysis further showed that murine cross-species correlations aligned primarily along a common mRNA axis; positive correlations with each murine model tended to occur at positive PC1 values and negative correlations at negative PC1 values (Supplemental Figure 7A-D). Prominent total-RNA loading features (centered, unscaled) included myeloid and innate-response genes, including *CD177, SIGLEC14, DAAM2*, and *DEFA3* (Supplemental Figure 7A).^38^

Because the human cohort was not stratified by adjudicated AKI etiology, these correlations were not intended to assign individual participants to discrete IRI-like, CLP-like, or LPS-like categories (Figure 7B). Rather, the analysis demonstrates that experimentally defined murine injury programs are represented to varying degrees within the heterogeneous molecular background of human AKI.

## Discussion

In this work, we defined distinct kidney and blood transcriptomic programs across murine IRI, CLP, and LPS injury models and leveraged these experimentally controlled states to interrogate molecular heterogeneity in human AKI. Experimental AKI etiologies were readily distinguishable in mice, although the underlying molecular features varied by both model and biological compartment. Kidney mRNA most clearly distinguished injury type: IRI was characterized by wound healing, cell adhesion, and metabolic suppression; CLP elicited the broadest response, combining inflammatory and secretory activation with suppression of mitochondrial, ciliary, and biosynthetic programs; and LPS induced a more restricted innate, antiviral, and interferon-associated signature. Blood retained etiology-specific information but primarily reflected systemic processes, with dominant inflammatory signaling in LPS, platelet and hemostatic programs in CLP, and comparatively modest changes in IRI; miRNA profiles were less able to distinguish distinct AKI etiologies.

When projected onto human AKI, the murine blood signatures mapped to a shared molecular continuum, rather than discrete murine-defined etiological groups. Importantly, this is consistent with the biological complexity of clinical AKI, where ischemia, infection, hemodynamic perturbation, comorbidities, and therapeutic interventions frequently coexist. Thus, the cross-species findings do not diminish the relevance of the murine etiology-specific signatures; rather, they illustrate how those controlled experimental programs are embedded within the more complex molecular architecture of human disease, reflective of the intersection of injury etiology, severity, and molecular stage, with repair and recovery programs evolving across this continuum.

Our murine findings extend prior work describing dynamic epithelial injury and repair states after IRI and temporally organized inflammatory responses during endotoxemia.^5; 8; 12; 13; 22; 27; 45; 46^ The wound-healing, adhesion, and metabolic programs observed in IRI are consistent with epithelial cell remodeling, whereas the innate immune and interferon response in LPS recapitulates key components of the host response previously described in endotoxemia.^5; 27^ CLP, however, was not simply a more severe form of LPS-induced injury. Its renal transcriptome combined secretory activation with suppression of mitochondrial, ciliary, and biosynthetic processes, while the corresponding blood profile was enriched for platelet activation, coagulation, and wound healing. These distinctions align with evidence that sepsis transcriptomes contain both shared host-response programs and context-specific modules.^47–49^ Most transcriptomic studies of AKI have focused on a single experimental model, molecular readout or tissue compartment.^3; 5; 12; 13; 22; 30; 31; 45; 46^ By directly comparing paired kidney and blood mRNA and miRNA across three AKI models at a common time point, our study fills this gap and demonstrates that experimentally distinct injuries capable of producing the same clinical AKI phenotype remain molecularly separable. These findings further caution against treating LPS and CLP as interchangeable models of sepsis-associated AKI and provide a framework for selecting the experimental model that best matches the biological pathway under investigation. This experimental framework next allowed us to ask whether these molecular programs are conserved within the substantially more heterogeneous setting of human AKI.

Several limitations warrant consideration. The human cross-species analysis should be interpreted as an assessment of molecular conservation rather than a direct etiologic classifier, as the human cohort was not stratified into purely ischemic, CLP-like, or endotoxemic AKI. Such classification may also be difficult in clinical AKI, where patients frequently experience overlapping insults and molecular subphenotypes that transcend conventional diagnostic categories.^2; 48; 50–52^ Circulating transcriptomes further integrate injury mechanisms with changes in cellular composition, systemic illness, comorbidities, and treatment; therefore, future studies should evaluate these signatures in larger, longitudinal cohorts with rigorous etiologic adjudication and blood-cell deconvolution, and determine whether combined etiology- and recovery-associated signatures improve prediction beyond KDIGO stage and conventional biomarkers.^53–55^ Additional limitations include the small murine sample size, use of young male mice, evaluation at a single 24-hour time point, and reliance on inferred rather than experimentally validated miRNA-mRNA interactions. These limitations are balanced by the paired kidney-blood design, parallel mRNA-miRNA profiling, direct comparison of three mechanistically distinct injuries, and cross-species analysis, which enabled examination of local renal injury and circulating host responses within a common framework. Importantly, signatures derived from these relatively mild experimental injuries were represented to varying degrees across human AKI and showed their strongest relationship with tubular casts, reflecting more severe pathologic injury and suggesting that milder experimental models may preserve etiology-specific biology relevant across the spectrum of human disease.

The translational value of these findings does not lie in classifying patients according to a single AKI etiology. Rather, experimentally derived signatures may serve as molecular reference programs that quantify the relative contribution of ischemic injury, inflammatory activation, metabolic dysfunction, coagulation, and tissue repair within an individual’s mixed clinical phenotype. Blood is particularly attractive for this purpose because it is serially accessible and, in our study, retained information related to injury etiology. A biomarker strategy integrating circulating mRNA, miRNA, and kidney injury proteins could ultimately identify active biological pathways, distinguish ongoing injury from adaptive repair, enrich clinical trials for patients most likely to respond to targeted interventions, and match treatment to mechanism rather than serum creatinine elevation alone.^51–53^ This cross-species framework provides a path toward biologically informed precision medicine for human AKI.

## Methods

### Sex as a biological variable

All animal experiments were performed in male C57BL/6J mice aged 8-12 weeks. Prior studies have documented sex-dependent responses in AKI and sepsis models; female mice have shown less renal functional and histologic injury after ischemia-reperfusion injury (IRI) than male mice, and in CLP and LPS models, sex-related effects vary with experimental conditions and the outcome measured.^56–58^ We therefore designed this study to compare injury etiologies in male mice because they exhibit a consistent injury phenotype; female mice should be evaluated directly to determine the relevance of these findings. In human studies, we included both males and females in the analysis.

### Animal Models

Male C57BL/6J mice (Jackson Laboratory, Bar Harbor, ME, USA) were 8-12 weeks of age (25-30 g). Each mouse was subjected to one model of AKI or relative experimental controls and was sacrificed 24 hours after onset of injury. All surgeries were performed under isoflurane anesthesia. IRI (n=5): Body temperature was continuously monitored with a rectal probe and maintained at 37°C with a temperature-controlled heating pad. After a midline incision and displacement of internal organs, the bilateral renal veins and pedicles were isolated and clamped with microvascular clips (Roboz Surgical Instrument Company, Gaithersburg, MD, USA) for 9 minutes. Ischemia was confirmed by a color change of the kidney from pink to pale. After 9 minutes, the clamps were removed, and reperfusion was verified by the return of the kidneys to a pink color; CLP (n=5, from which 4 had blood and kidney tissue available for sequencing analysis): a midline incision was performed to expose the cecum which was then externalized. The ileocecal valve was identified; then 75% of the mouse cecum was ligated and punctured once with a 25g needle, and a small amount of fecal material was extruded through the puncture site; the cecum was then returned to the abdominal cavity. LPS (n=3): Mice were subjected to a single dose of 2.5 mg/kg LPS (E.coli serotype O111:B4, Sigma-Aldrich, St. Louis, MO, USA) via tail vein injection (i.v.). No antibiotics were administered to any mice. Sham-operated control mice (n=3) underwent identical surgical procedures without cecal puncture or renal vasculature clamping. All mice that underwent surgery were resuscitated with normal saline subcutaneously (500 µL, s.q.) and post-operative analgesia with buprenorphine XR (3.5 mg/kg, s.q.). Controls for LPS (n=3) were injected with an equivalent amount of vehicle (normal saline, i.v.).

### Tissue collection and fixation

Bilateral kidneys were obtained, capsules removed and cut transversely, and 300 µL of blood was obtained via intracardiac puncture at sacrifice. Kidneys were fixed in 4% formalin, transferred to 70% ethanol for paraffin embedding, and sectioned at 5 µm and stained with hematoxylin and eosin (H&E) for histologic analysis, or frozen in liquid nitrogen and stored at −80°C for real-time PCR or total RNA/miRNAseq. A total of 300 µL of blood was collected and divided into 200 µL for downstream serum creatinine analysis, and a 100 µL aliquot preserved in RNAprotect Animal Blood Tubes (100 µL, Qiagen, Germantown, MD, USA) for downstream miRNA and total RNAseq.

### Histologic examination

Histologic changes in the renal cortex and outer stripe of the outer medulla were evaluated separately in five non-overlapping fields per compartment per representative mouse (n=1, H&E stain, 20× magnification) by an investigator blinded to experimental group. We applied the same field-selection criteria and magnification to all animals. Tubular epithelial injury was assessed semi-quantitatively based on the percentage of unique tubules exhibiting at least one of the following features: loss of brush border, tubular dilation, cast formation, or epithelial necrosis/lysis or sloughing. We counted tubules exhibiting one or more injury features once, regardless of the number of features present. Injury was scored as follows: 0, no injured tubules; 1, <10%; 2, 10–25%; 3, 26–50%; 4, 51–75%; and 5, >75% injured tubules. Scores from the five fields within each compartment were averaged to obtain one cortical and one outer-medullary injury score per mouse. Digital images were viewed and analyzed using Fiji (ImageJ).^59; 60^

### Measurement of Serum Creatinine (sCr)

Whole blood samples were centrifuged at 3,000 g for 10 minutes at 4°C to separate serum for analysis of sCr. The measurement of sCr was provided by the University of Alabama at Birmingham/UCSD O’Brien Center Core for Acute Kidney Injury Research (NIH P30DK079337) using isotope dilution liquid chromatography-tandem mass spectrometry.

### Total RNA Isolation

<u>Murine Tissues:</u> For the murine samples, half of each frozen kidney was homogenized in QIAzol Lysis Reagent (Qiagen), and whole blood was collected in RNAprotect Animal Blood Tubes. Total RNA was isolated from kidney and blood samples using the miRNeasy Kit (Qiagen, Germantown, Maryland, USA), including DNase treatment, according to the manufacturer’s instructions. RNA concentration and purity were assessed using a NanoPhotometer NP80 UV-Vis spectrophotometer (Implen, Westlake Village, CA). <u>Human blood:</u> Whole blood samples from 48 participants enrolled in the KPMP AKI cohort were obtained through an approved KPMP ancillary study (*Transcriptomic diagnosis and staging of human acute kidney injury*, AS5).^61^ Human blood collection, total RNA isolation, and RNA quality control were performed by the KPMP according to KPMP standard protocols, and RNA isolates were distributed by the University of Michigan Central Biorepository. Total RNA was then sequenced at Indiana University as described below. Human whole-blood miRNA libraries were also sequenced, but the resulting reads were not processed or analyzed for this study; all human analyses reported here used total RNA-seq data only.

### Quantitative Real-Time PCR (qRT-PCR)

cDNA was synthesized using the qScript cDNA Synthesis Kit (Quantabio, Beverly, MA) according to the manufacturer’s instructions. Expression of *Havcr1* and *Lcn2* was quantified by probe-based multiplex qRT-PCR using PrimeTime Gene Expression Master Mix (Integrated DNA Technologies, Coralville, IA) on a QuantStudio 3 Real-Time PCR System (Applied Biosystems/Thermo Fisher Scientific, Foster City, CA). Custom murine-specific TaqMan primers and probes for *Gapdh* (TAMRA), *Havcr1* (FAM), and *Lcn2* (VIC) were used for multiplex real-time PCR, with ROX serving as the passive reference dye (Thermo Fisher Scientific, Waltham, MA). Reactions were performed in technical triplicate in a final volume of 20 µL. PCR cycling consisted of polymerase activation at 95°C for 3 minutes followed by 40 cycles of denaturation at 95°C for 5 seconds and annealing/extension at 60°C for 30 seconds, with fluorescence acquisition during the annealing/extension step. Mean Ct values from technical replicates were normalized to *Gapdh*, and relative expression was calculated using the ΔΔCt method with the respective sham or saline control group as the experimental reference control and expressed as log2 fold change relative to control. *total RNAseq (murine and human)*: Total RNA samples were first evaluated for their quantity and quality using Agilent TapeStation. One hundred nanograms of total RNA was used for library preparation with the Illumina Stranded Total RNA Prep, Ligation with Ribo-Zero Plus kit (Illumina, San Diego, CA), following the manufacturer’s instructions. Globin RNA removal was integrated into the protocol for whole blood samples. Each resulting uniquely dual-indexed library was quantified and quality assessed by Qubit and Agilent TapeStation, and multiple libraries were pooled in equal molarity. The pooled libraries were sequenced with 2×150bp paired-end configuration on an Illumina NovaSeq X PLUS sequencer.

### miRNA sequencing (murine and human)

The same samples used for the total RNA-seq were utilized for the miRNA library preparation. 10 ng of total RNA was used for the miRNA libraries construction with the QIAseq miRNA Library Kit for Illumina (Cat. No. 331505; Qiagen, Germantown, MD). Each resulting indexed library was quantified and its quality assessed by using Qubit fluorometric quantification (Thermo Fisher Scientific, Waltham, MA) and Agilent TapeStation. The final libraries were sequenced with 76 bp single-end read configuration on an Illumina NovaSeq X PLUS sequencer (Illumina, San Diego, CA). 12-20 million reads were targeted for each sample.

### Alignment of sequence data and QC: miRNA

The de-multiplexed raw sequence reads were uploaded to the Qiagen GeneGlobe Data Analysis Center for quality control, alignment and expression quantification and reads mapped to different databases (miRbase mature, miRbase hairpin, noncoding RNA, mRNA and otherRNA using bowtie v1.2. ^62^ Read and UMI counts for each small RNA category calculated for each sample. MiRbase v21 was used for miRNA. *mRNA*: The sequencing data was assessed for quality using FastQC, then mapped to reference genomes: mm10 for mice, and hg38 for humans, gene counts quantified and further standard RNAseq quality control assessments to generate raw counts table for each miRNA and mRNA seq data, respectively.

### Murine Data Analysis

All bioinformatic and statistical analyses were performed in R version 4.4.1, with the complete computational environment and package versions provided in Supplemental Text File. Bulk RNA-seq analyses included CLP (n=4), IRI (n=5), LPS (n=3), sham (n=3), and saline controls (n=3); five mice underwent CLP, of which four had samples available for transcriptomic analysis. Gene symbols were updated to current MGI nomenclature using org.Mm.eg.db, and only mature mouse miRNAs were retained (mmu-miR, mmu-let). mRNA and miRNA count matrices were analyzed using edgeR.^63^ Lowly expressed features were removed using filterByExpr() (min.count=1, min.total.count=3), libraries were normalized by the trimmed mean of M-values (TMM) method, and normalized log2 counts per million (CPM) were calculated with a prior count of 2. Differential expression was assessed using a cell-means design (∼0 + group) and quasi-likelihood models fitted with glmQLFit(). IRI and CLP were compared with sham controls and LPS with saline controls. Differentially expressed features were identified using glmTreat() with FDR <0.05 and absolute fold change ≥1.2. PCA was performed on scaled log2 CPM matrices, and features contributing most strongly to principal components were identified from PCA loadings. Differentially expressed features were intersected separately by direction to identify model-specific, AKI-shared, and CLP/LPS-shared responses. Heatmaps were generated from log2 CPM values, with treatment-specific and intersection features ranked by absolute fold change and standardized to z-scores where indicated. GO Biological Process enrichment was performed using clusterProfiler enrichGO() with Benjamini-Hochberg correction (P<0.05, q<0.20), using all genes tested in the corresponding contrast as the background.

Integrated miRNA-mRNA analysis was performed using mirTarRnaSeq and miRDB v6.0 interactions with prediction scores ≥80.^32; 64^ mRNAs with glmQLFTest P>0.05 were excluded, and miRNA-mRNA interaction scores were calculated from the divergence in fold change between each pair. A null distribution was generated from 100 samples of 1,000 randomly selected miRNA-mRNA pairs, and candidate interactions exceeding the 95th percentile were retained. Interactions were further restricted to miRDB-supported pairs in which the miRNA was differentially expressed at FDR <0.05. For each AKI model, a representative treatment-specific upregulated kidney miRNA was selected based on fold change and availability of predicted target interactions. Target-gene modules were evaluated by GO enrichment and subsequently projected onto single-cell kidney transcriptomic data.

Publicly available murine kidney scRNA-seq datasets were analyzed using Seurat for IRI (GSE171639; wild-type 30-min ischemia), CLP (GSE298053; GSM9006031), and LPS (GSE151658; LPS 4-hour time point).^5; 46; 65; 66^ Cells were quality-filtered according to dataset-specific RNA feature and molecule-count thresholds and <50% mitochondrial and <20% ribosomal transcripts. Datasets were independently normalized using SCTransform() with 3,000 variable features and mitochondrial percentage regressed out, followed by PCA. Published kidney cell-type annotations from the LPS dataset were transferred to IRI and CLP using the top 30 principal components. Datasets were then merged, renormalized, and integrated using canonical correlation analysis (CCA), and UMAP was generated from the top 30 integrated dimensions. Predicted target genes for representative miRNAs were scored at the single-cell level using AddModuleScore(), and target networks were annotated according to differential expression in the corresponding bulk kidney RNA-seq analysis.

### Human Data Analysis

Human whole-blood total RNA sequencing data were analyzed from 48 KPMP participants enrolled in the AKI cohort using baseline samples. Gene counts were analyzed using edgeR, with lowly expressed genes removed using filterByExpr() (min.count=1, min.total.count=3), retaining 15,681 genes. Libraries were normalized using the trimmed mean of M-values (TMM) method, and log2 counts per million (CPM) were calculated with a prior count of 2. Participant sex was regressed from the log2 CPM matrix using limma::removeBatchEffect(). Human-mouse orthologues were obtained from Ensembl (release 116) using biomaRt, retaining high-confidence one-to-one orthologues, yielding 15,451 gene pairs; 10,568 genes present in both the filtered human dataset and all three murine blood contrasts were used for cross-species analysis.^67; 68^ Each human participant was compared with each mouse AKI model by rank correlation. On the mouse side, we used the whole-blood totalRNA log2 fold-changes for IRI vs Sham, CLP vs Sham, and LPS vs Saline from the edgeR models described for the mouse bulk RNA-seq; this included every tested gene rather than only the DE genes. On the human side, we centered each gene’s sex-adjusted log2 CPM by subtracting its mean across all 48 participants, so each value reflects deviation from the cohort mean, comparable to a mouse fold-change. The Spearman correlation between each participant’s centered values and each mouse contrast’s fold-changes was computed over the working gene set, and each participant’s mean mouse correlation was defined as the mean of their IRI, CLP, and LPS correlations.

PCA was performed on the sex-adjusted human log2 CPM matrix using all 15,681 filtered genes, with both centered-only and centered-and-scaled analyses performed. The top 10 contributors to PC1 and PC2 were identified by loading-vector magnitude, and participants were projected onto PC1 and PC2 for comparison with biopsy tubular casts and model-specific murine correlation scores.

### Human Clinical Data Collection and AKI Staging

Associated clinical, demographic, and adjudicated data were obtained from the KPMP database. In baseline visit data: AKI severity was graded according to the Kidney Disease: Improving Global Outcomes (KDIGO) clinical practice guidelines.^1^ Participants were grouped based on their peak serum creatinine (sCr) relative to baseline values: Stage 1, Stage 2, Stage 3, or No AKI. Baseline sCr was determined using documented historical actual values when available; in cases where historical data were missing, baseline sCr was calculated via standard KPMP imputation protocols.

### Human Kidney Tissue:Histopathologic Assessment

Histologic evaluation was performed on baseline renal biopsy specimens collected at the time of KPMP enrollment. The KPMP central pathology adjudication committee systematically assessed and quantified the visual presence or absence of histopathologic casts (including tubular, granular, or cellular casts). These histopathologic findings were subsequently cross-referenced with the corresponding clinical demographic profiles and adjudicated KDIGO AKI severity stages for downstream statistical analysis.

### Statistical analysis

Murine mRNA and miRNA sequencing counts were analyzed using edgeR. Lowly expressed features were filtered using filterByExpr() (minimum count = 1; minimum total count = 3), and libraries were normalized using the trimmed mean of M-values method.

Differential expression was assessed using quasi-likelihood models fitted with glmQLFit() and tested with glmTreat(). Features with a false discovery rate below 0.05 and an absolute fold change of at least 1.2 were considered differentially expressed. Gene Ontology Biological Process enrichment was performed using clusterProfiler, with Benjamini-Hochberg correction; terms with P < 0.05 and q < 0.20 were considered enriched. Human-mouse expression associations were assessed by Spearman rank correlation across 10,568 high-confidence one- to-one orthologs. qRT-PCR, serum creatinine, and histologic injury scores are presented descriptively; no inferential statistical tests were performed for these outcomes.

### Study approval

All animal protocols were approved by the Indiana University Institutional Animal Care and Use Committee and performed in accordance with the National Institutes of Health Guide for the Care and Use of Laboratory Animals. Human blood samples and clinical datasets were obtained through the Kidney Precision Medicine Project (KPMP) consortium through an approved ancillary study. All participants provided written informed consent, and all study activities were reviewed and approved under a protocol by the KPMP single IRB of the University of Washington (IRB #20190213).

### Data Availability

Murine data in this publication have been deposited in the National Center for Biotechnology Information’s Gene Expression Omnibus database (https://www.ncbi.nlm.nih.gov/geo) and are accessible through accession GSE349874 (totalRNAseq-murine blood and kidney), GSE349746 (miRNAseq-murine blood and kidney). Additional murine data previously published are accessible via: GSE171639, GSE298053; GSM9006031, GSE151658. The human results here are in part based upon data generated by the Kidney Precision Medicine Project, accessed 7/2026: https://www.kpmp.org. Human blood sequencing data will be released from the Kidney Precision Medicine Project database upon acceptance of this study and made available according to their data use agreement.

## Supporting information

KPMP Consortium List

## Author contributions

DJ, PD, TH, MTE conceived the study and designed the research. DJ performed the experiments. DJ acquired the data. DJ, TWMC, AS analyzed the data. DJ and KPMP provided clinical data, samples, or other resources. All authors drafted and revised the manuscript. All authors reviewed and approved the final manuscript.

## Conflict of interest

The authors have declared that no conflict of interest exists.

## Funding statement

DJ is supported by the NIH/National Institute of Diabetes and Digestive and Kidney Diseases (NIDDK) (K08DK139369), NIH/National Center for Advancing Translational Sciences-Clinical and Translational Science Awards (UL1TR002529; S. Moe and S. Wiehe, co-Principal Investigators), a research grant from Dialysis Clinic, Inc., grant 2021258 from the Doris Duke Charitable Foundation through the COVID-19 Fund to Retain Clinical Scientists collaborative grant program; and grant 62288 from the John Templeton Foundation and the Department of Medicine, Indiana University School of Medicine; PD is supported by NIH/NIDDK (R01 DK107623; U01 DK114923), MTE is supported by U01 DK114923. TH is supported by VA Merit BX002901. This research was supported in part by Shared University Research grants from IBM, Inc., to Indiana University and the Lilly Endowment, Inc., through its support for the Indiana University Pervasive Technology Institute. The Kidney Precision Medicine Project (KPMP) is supported by the National Institute of Diabetes and Digestive and Kidney Diseases (NIDDK) through the following grants: U01DK133081, U01DK133091, U01DK133092, U01DK133093, U01DK133095, U01DK133097, U01DK114866, U01DK114908, U01DK133090, U01DK133113, U01DK133766, U01DK133768, U01DK114907, U01DK114920, U01DK114923, U01DK114933, U24DK114886, UH3DK114926, UH3DK114861, UH3DK114915, UH3DK114937, 1U01DK144965-01, 1U01DK144994-01.

## Acknowledgements

We gratefully acknowledge the essential contributions of our patient participants and the support of the American public through their tax dollars. The opinions expressed in this publication are those of the author(s) and do not necessarily reflect the view of the John Templeton Foundation. The content is solely the responsibility of the authors and does not necessarily represent the official views of the NIH. The authors acknowledge the University of Michigan Medical School Central Biorepository (RRID:SCR_026845) for providing biospecimen storage, management, and distribution services in support of the research reported in this publication.

## AI disclosure

The authors used ChatGPTEdu (−5.6 Sol) and Claude Opus 5.5 to assist with writing and refining code, which were then reviewed and verified by the authors (dates of use: February 2026-October 2026). AI tools were not used to generate or modify research data or data images. The authors assume full responsibility for ensuring the integrity and accuracy of data output presented in this article.

---

**Supplemental Figure 1.**
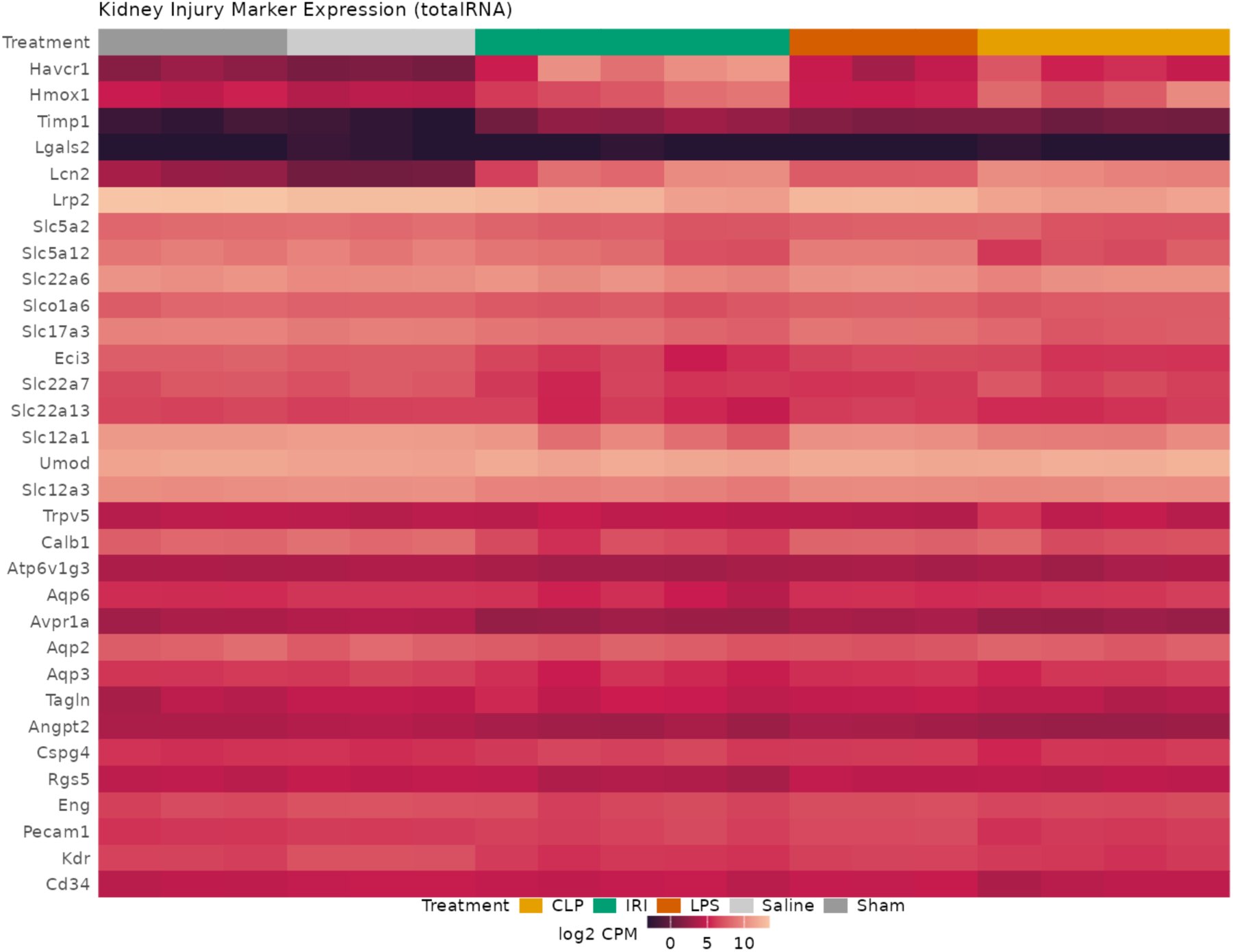
Kidney expression of canonical renal cell and acute tubular injury markers across murine AKI models. Heatmap showing expression of selected canonical renal cell markers and acute tubular injury markers across kidney samples, ordered by treatment group. Expression is shown as log2 counts per million (CPM). IRI, ischemia-reperfusion injury; CLP, cecal ligation and puncture; LPS, lipopolysaccharide; CPM, counts per million.

**Supplemental Figure 2.**
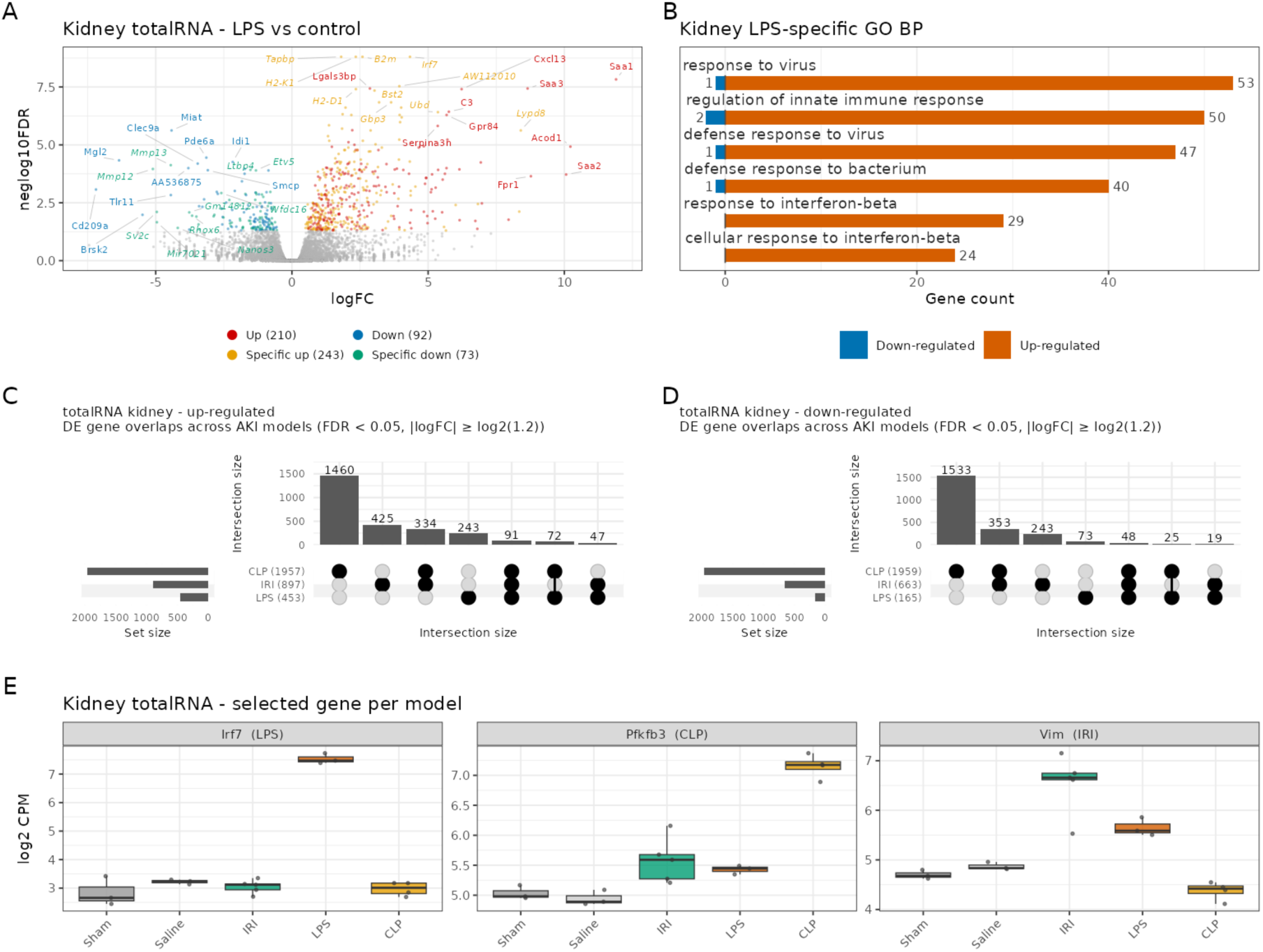
Shared and model-specific kidney transcriptional responses across murine AKI models. (A) Volcano plot of kidney total RNA differential expression in LPS versus control. (B) GO Biological Process terms enriched among LPS-specific kidney total RNA DEGs. (C and D) UpSet plots of upregulated (C) and downregulated (D) kidney total RNA DEGs shared across AKI models. (E) Expression of representative model-specific genes (*Vim*, IRI; *Pfkfb3*, CLP; *Irf7*, LPS) across treatment groups. Expression is shown as log2 CPM; boxes indicate median and interquartile range, and points represent individual samples. DEG cutoffs as previously described in Figure 2 legend. IRI, ischemia-reperfusion injury; CLP, cecal ligation and puncture; LPS, lipopolysaccharide; GO, Gene Ontology; DEG, differentially expressed gene; CPM, counts per million.

**Supplemental Figure 3.**
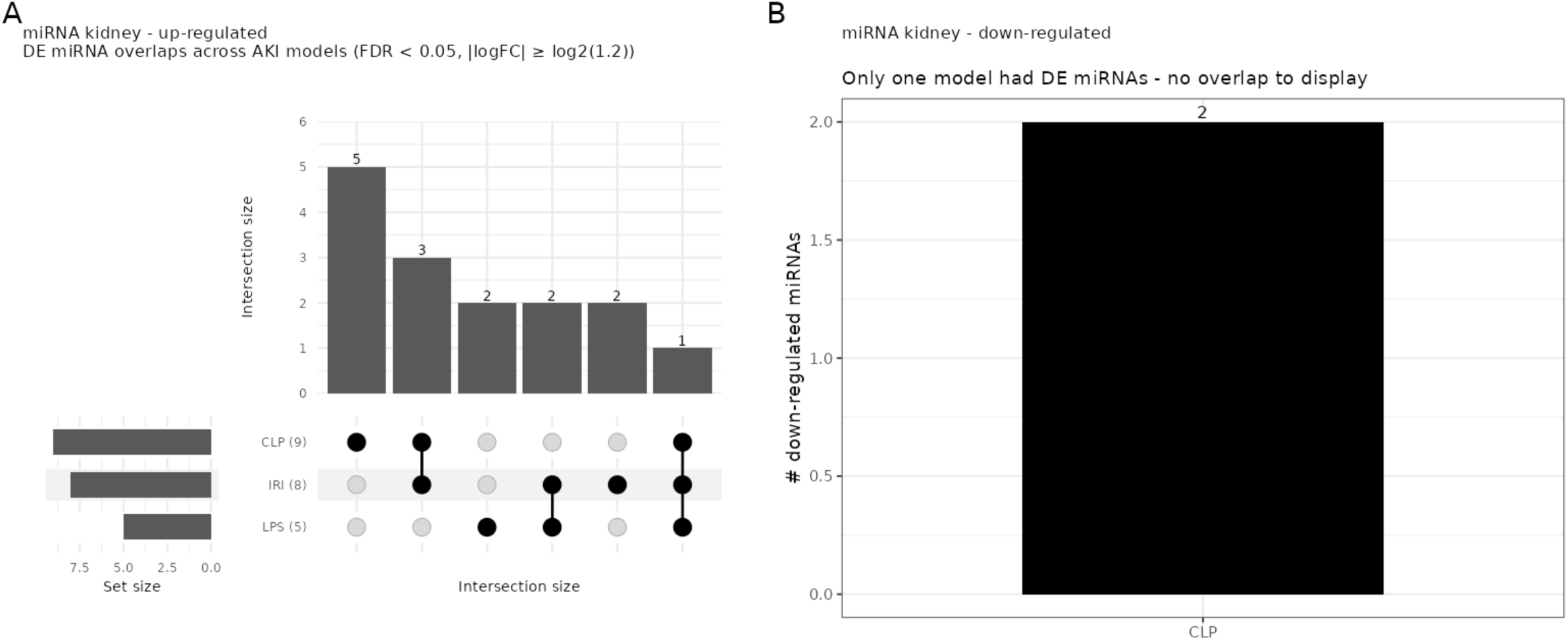
Overlap of differentially expressed kidney miRNAs across murine AKI models. (A) UpSet plot showing intersections of upregulated kidney miRNAs across IRI, CLP, and LPS models. Set sizes are indicated in parentheses. (B) Number of downregulated kidney miRNAs by AKI model; downregulated miRNAs were identified only in CLP, precluding intersection analysis. Differentially expressed miRNAs were defined as previously described in Figure 3 legend. IRI, ischemia-reperfusion injury; CLP, cecal ligation and puncture; LPS, lipopolysaccharide; FDR, false discovery rate.

**Supplemental Figure 4.**
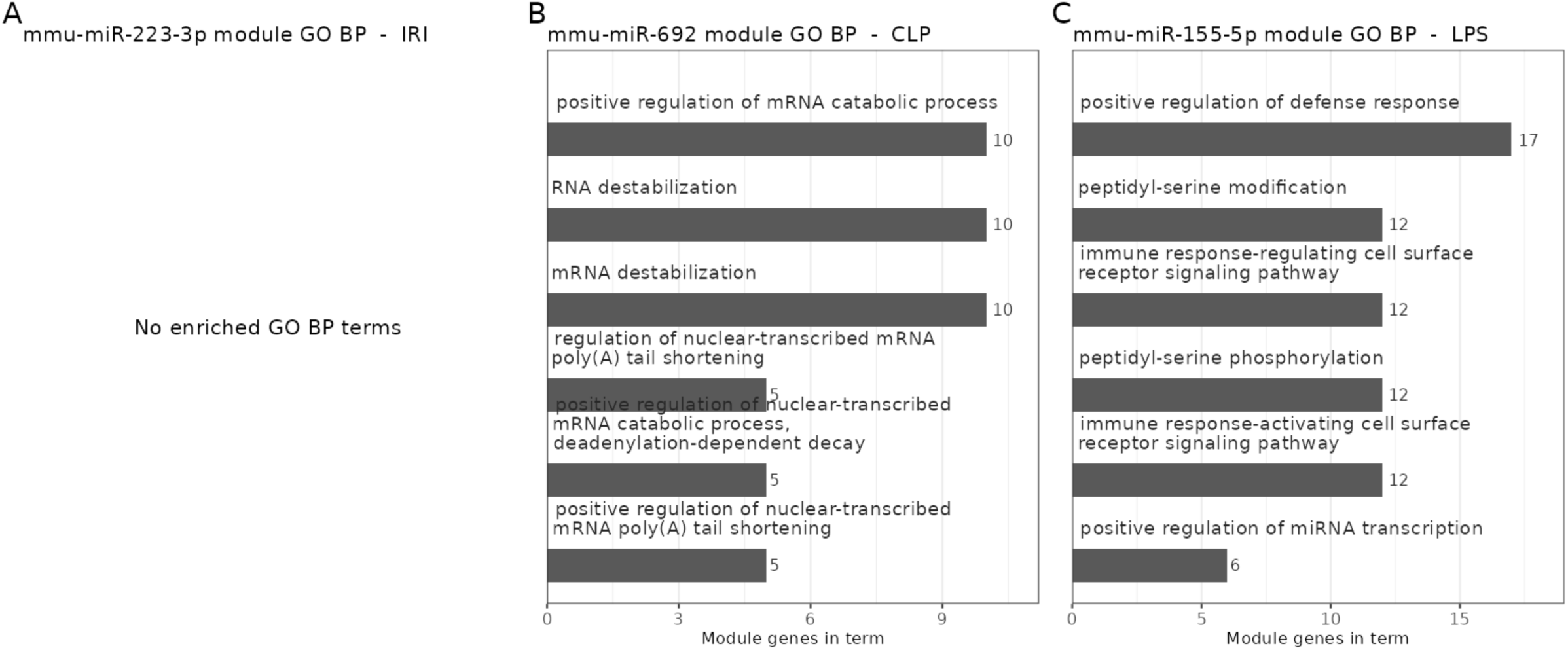
Biological processes enriched among predicted miRNA target modules in experimental AKI. Gene Ontology (GO) Biological Process enrichment of predicted target modules for mmu-miR-223-3p in IRI (A), mmu-miR-692 in CLP (B), and mmu-miR-155-5p in LPS (C). Up to 6 enriched terms are shown; bars indicate the number of module genes annotated to each term. No significantly enriched GO Biological Process terms were identified for the mmu-miR-223-3p target module (A). Enrichment was performed using the complete predicted target modules, with all genes detected in the integrated single-cell dataset as the background. IRI, ischemia-reperfusion injury; CLP, cecal ligation and puncture; LPS, lipopolysaccharide; GO, Gene Ontology.

**Supplemental Figure 5.**
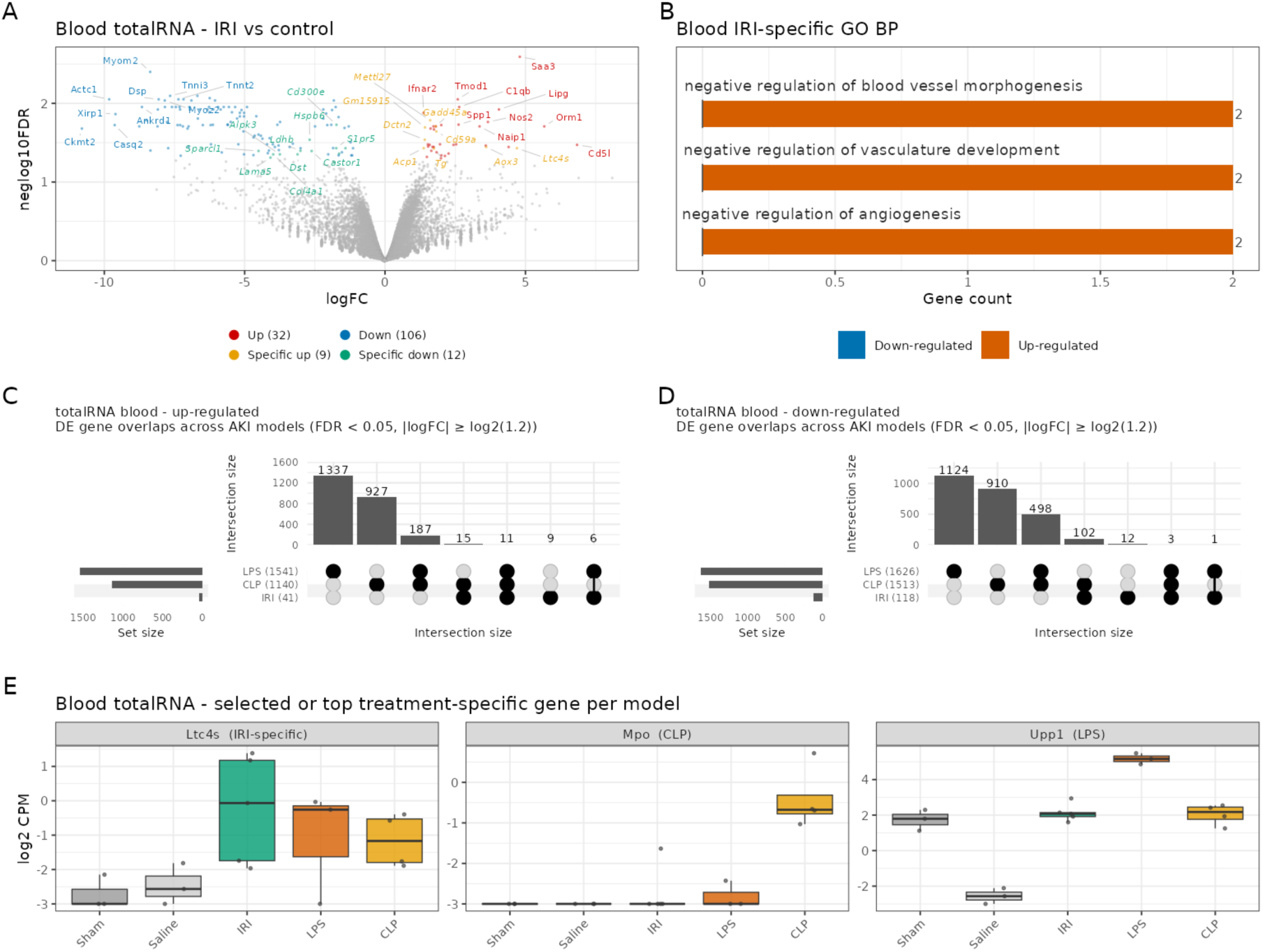
Shared and model-specific blood mRNA responses across murine AKI models. (A) Volcano plot of blood total RNA differential expression in IRI versus control. (B) GO Biological Process terms enriched among IRI-specific blood total RNA DEGs. (C and D) UpSet plots showing intersections of upregulated (C) and downregulated (D) blood total RNA DEGs across IRI, CLP, and LPS models. (E) Expression of representative treatment-specific genes (*Ltc4s*, IRI; *Mpo*, CLP; *Upp1*, LPS) across treatment groups. Expression is shown as log2 CPM; boxes indicate median and interquartile range, and points represent individual samples. DEGs were defined as previously described in the Figure 2 legend. IRI, ischemia-reperfusion injury; CLP, cecal ligation and puncture; LPS, lipopolysaccharide; GO, Gene Ontology; DEG, differentially expressed gene; CPM, counts per million.

**Supplemental Figure 6.**
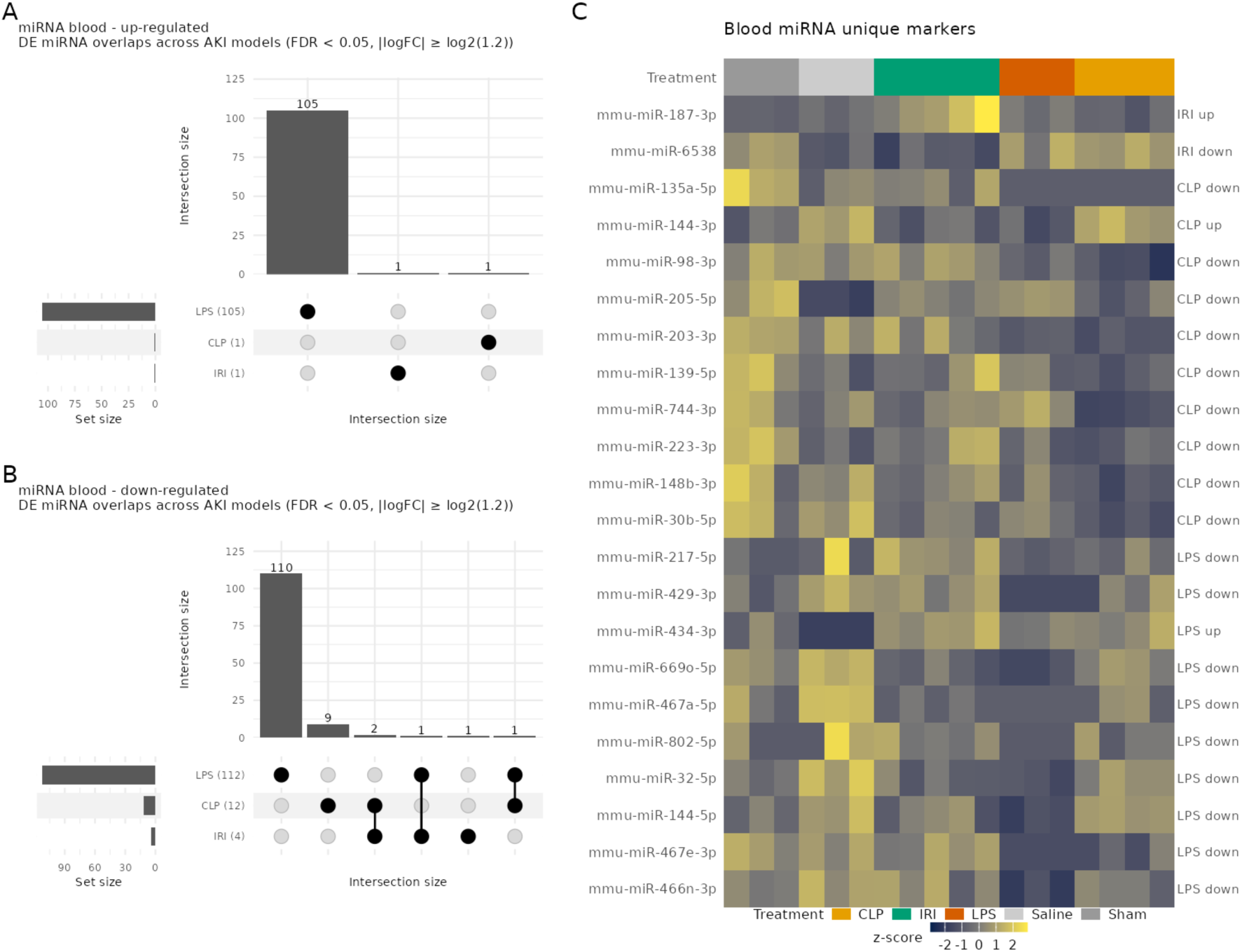
Shared and model-specific blood miRNA expression across experimental AKI models. (A and B) UpSet plots showing intersections of upregulated (A) and downregulated (B) differentially expressed blood miRNA across IRI, CLP, and LPS models. Set sizes are indicated in parentheses. (C) Heatmap of treatment-specific blood miRNA, ordered by treatment group and shown as row-scaled z scores, ranked by absolute fold change; all available markers are shown when fewer than 10 were identified. DE miRNA cutoffs previously defined in Figure 3 legend. IRI, ischemia-reperfusion injury; CLP, cecal ligation and puncture; LPS, lipopolysaccharide; DE, differentially expressed; FDR, false discovery rate.

**Supplemental Figure 7.**
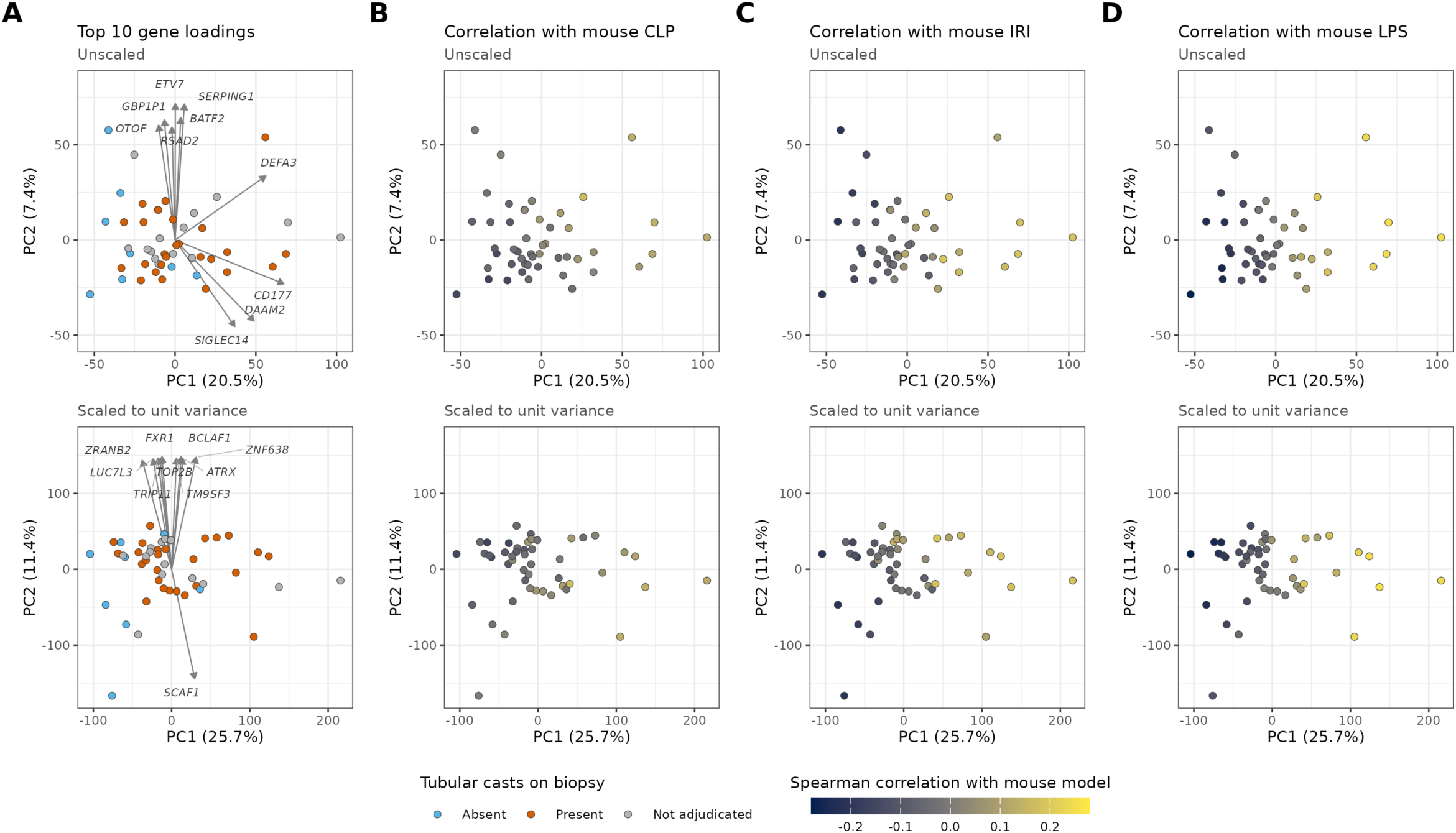
Murine AKI signatures align with a shared transcriptomic axis in human AKI. (A) PCA of whole-blood total RNA from 48 KPMP participants enrolled in the AKI cohort, colored by tubular casts on biopsy (absent, n=8; present, n=26; not adjudicated, n=14). Arrows indicate the 10 genes with the largest PC1-PC2 loadings. (B-D) Samples colored by Spearman correlation with murine blood signatures for (B) CLP, (C) IRI, and (D) LPS. PCA was performed on sex-adjusted log2 CPM across 15,681 filtered genes. Top panels show centered, unscaled PCA (PC1, 20.5%; PC2, 7.4%); bottom panels show PCA after scaling genes to unit variance (PC1, 25.7%; PC2, 11.4%). Correlations with all three murine models varied predominantly along PC1 and were similar with and without scaling. Each point represents one participant.

## Notes

### Competing Interest Statement

The authors have declared no competing interest.

